# Industrial bees: when agricultural intensification doesn’t impact local disease prevalence

**DOI:** 10.1101/428656

**Authors:** Lewis J. Bartlett, Carly Rozins, Berry J. Brosi, Keith S. Delaplane, Jacobus C. de Roode, Andrew White, Lena Wilfert, Michael Boots

## Abstract

1. Although it is generally thought that the intensification of farming will result in higher disease prevalences there is little specific modelling testing this idea. We build multi-colony models to inform how ‘apicultural intensification’ is predicted to impact honeybee pathogen epidemiology at the apiary scale.
2. Counter to the prevailing view, our models predict that intensification, captured though increased population sizes, changes in population network structure, and increased between-colony transmission, is likely to have little effect on disease prevalence within an apiary.
3. The greatest impacts of intensification are found for diseases with relatively low R_0_ (basic reproduction number), however, such diseases cause little overall disease prevalence and therefore the impacts of intensification are minor. Furthermore, the smallest impacts of intensification are found for diseases with high R_0_ values, which we argue are typical of important honeybee diseases.
4. *Policy Implications:* Our findings highlight a lack of support for the hypothesis that current and ongoing intensification leads to notably higher disease prevalences. More broadly, our work demonstrates the need for informative models of agricultural systems and management practices in order to understand the implications of management changes on diseases.

## Introduction

Infectious diseases exact tolls on agricultural sustainability (Brijnath, Butler, & McMichael, 2014) and profitability (James, 1981). A key question is how agricultural intensification and novel agricultural practices impact the emergence and epidemiology of infectious disease (Cressler, McLeod, Rozins, Hoogen, & Day, 2016; Gandon, Hochberg, Holt, & Day, 2013). A general assumption is that intensification increases vulnerability to severe disease outbreaks (Jones et al., 2013; Kennedy et al., 2016; Mennerat, Nilsen, Ebert, & Skorping, 2010), but there is relatively little empirical data we can use to understand how different agricultural approaches influence infectious disease prevalences, epidemiological theory is therefore a useful alternative (Atkins et al., 2013; Rozins & Day, 2016). Here we build specific models of apiary-level intensification in commercially farmed honeybees to examine the impact of industrial-scale management practices on honeybee infectious disease prevalence.

Honeybee health and the apicultural industry are under threat from a variety of pressures (Ghazoul, 2005; vanEngelsdorp & Meixner, 2010), including parasites and pathogens (Budge et al., 2015; De la Rúa, Jaffé, Dall’Olio, Muñoz, & Serrano, 2009; Potts et al., 2010). There is a growing body of literature documenting the damage that emerging or re-emerging diseases (Wilfert et al., 2016) are causing in apiculture (Jacques et al., 2017; Kielmanowicz et al., 2015) and native pollinators (Cohen, Quistberg, Philpott, & DeGrandi-Hoffman, 2017; Fürst, McMahon, Osborne, Paxton, & Brown, 2014; Graystock, Blane, McFrederick, Goulson, & Hughes, 2016; Manley, Boots, & Wilfert, 2015; McMahon et al., 2015; McMahon, Wilfert, Paxton, & Brown, 2018). Evidence exists supporting a link between the risk of these diseases and specific apicultural practices (Giacobino et al., 2014; Mõtus, Raie, Orro, Chauzat, & Viltrop, 2016; Pacini et al., 2016). However, the evidence is geographically limited, lacking in mechanistic underpinning, or contradictory even within this small collection of studies. It is therefore critical that we learn how different apicultural practices impact disease outcomes (Brosi, Delaplane, Boots, & Roode, 2017). The need for an epidemiological framing of honeybee diseases has been frequently discussed (Brosi et al., 2017; Fries & Camazine, 2001) in both empirical (van Engelsdorp et al., 2013) and modelling (Becher, Osborne, Thorbek, Kennedy, & Grimm, 2013) studies, but we lack a modelling framework for disease ecology in honeybees at a scale larger than a single colony.

Honeybees are typically managed in apiaries, which are associated colonies placed together for beekeeping convenience at a single site. Pathogen dynamics at the apiary level are determined both by pathogen transmission within and between colonies. Intensification of apiculture changes apiary ecology in a number of ways, all potentially relevant to disease (Brosi et al., 2017). In particular, increasing the number of colonies and changing the arrangement of those colonies influences epidemiology through changes in both the size and network structure of the population. They both may also increase the rate at which transmission between colonies occurs via more frequent ‘drifting’ of honeybees (Free, 1958; Neumann, Radloff, Pirk, & Hepburn, 2003). Drift is a key mechanism of between-colony pathogen transmission (Goodwin, Perry, & Houten, 1994; Roetschi, Berthoud, Kuhn, & Imdorf, 2008) and has been invoked as an explanatory mechanism accounting for higher disease prevalences in larger apiaries (Mõtus et al., 2016).

The intensification of agricultural systems generally means larger, denser population sizes and greater pathogen transmissibility at the local and landscape scale. To understand these effects in honeybees we build multi-colony models to examine how apicultural intensification is predicted to impact honeybee pathogen epidemiology. We examine the epidemiological consequences of increasing the number of colonies within an apiary, changing colony configurations, and increasing between-colony pathogen transmission.

## Methods

We combine mathematical models and agent-based model (ABM) simulations to make predictions on how intensification affects disease risk, spread, and endemic prevalence within an apiary. The key to our approach is that we capture pathogen transmission both within and between colonies.

We generalise colony arrangements to three unique configurations: array, circular and lattice (Fig. 1). We restrict between-colony pathogen transmission to nearest neighbours (see discussion), those in closest proximity to each other (connected by an arrow in Fig. 2). Between-colony transmission is always assumed to be at a lower rate than within colony transmission. The mathematical model allows us to obtain tractable analytical results while the ABM simulations allow us to model disease at the level of the individual bee and consider stochastic effects.

**Figure 1.**
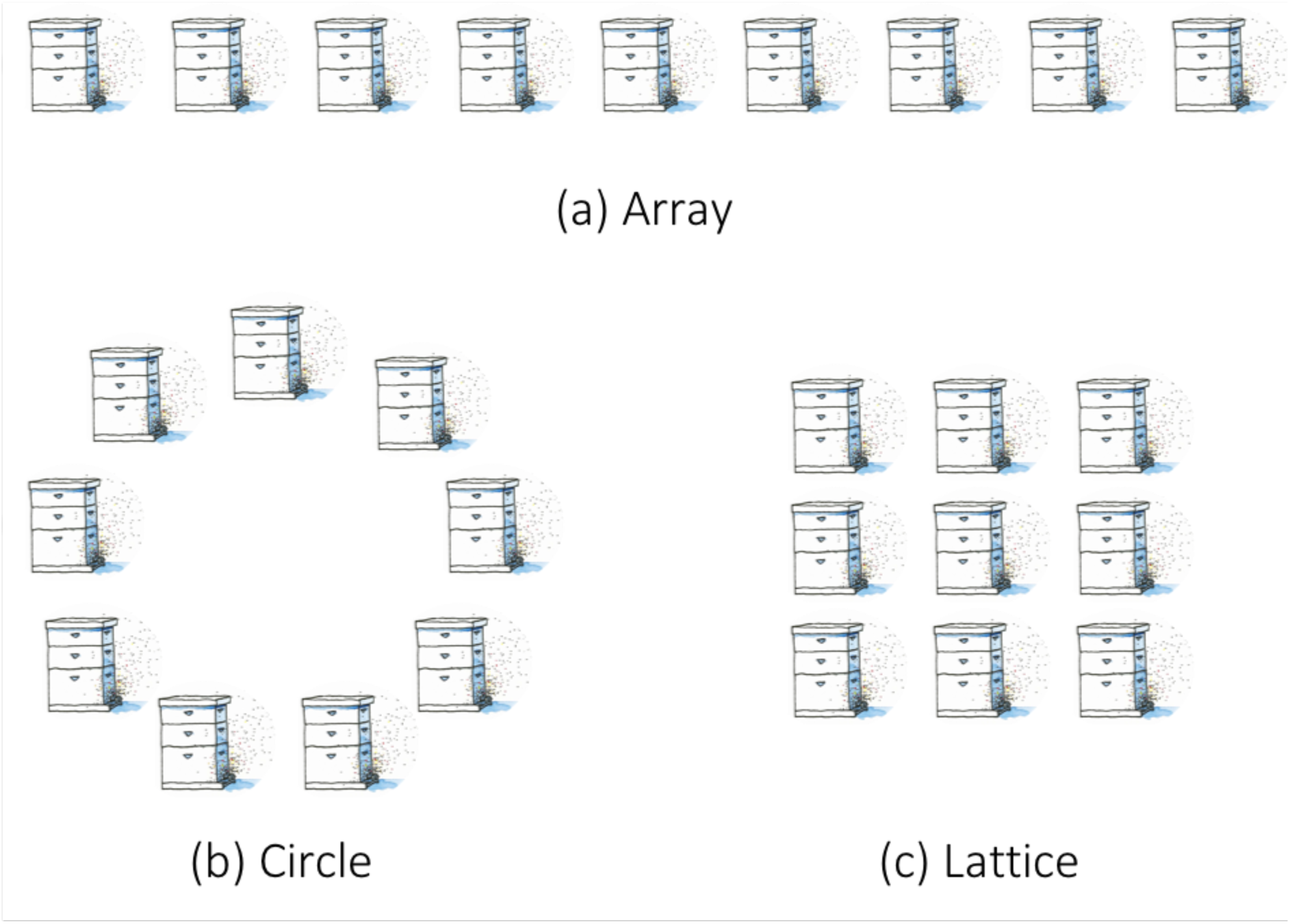
Colony configurations, demonstrated for apiaries with nine colonies.

**Figure 2.**
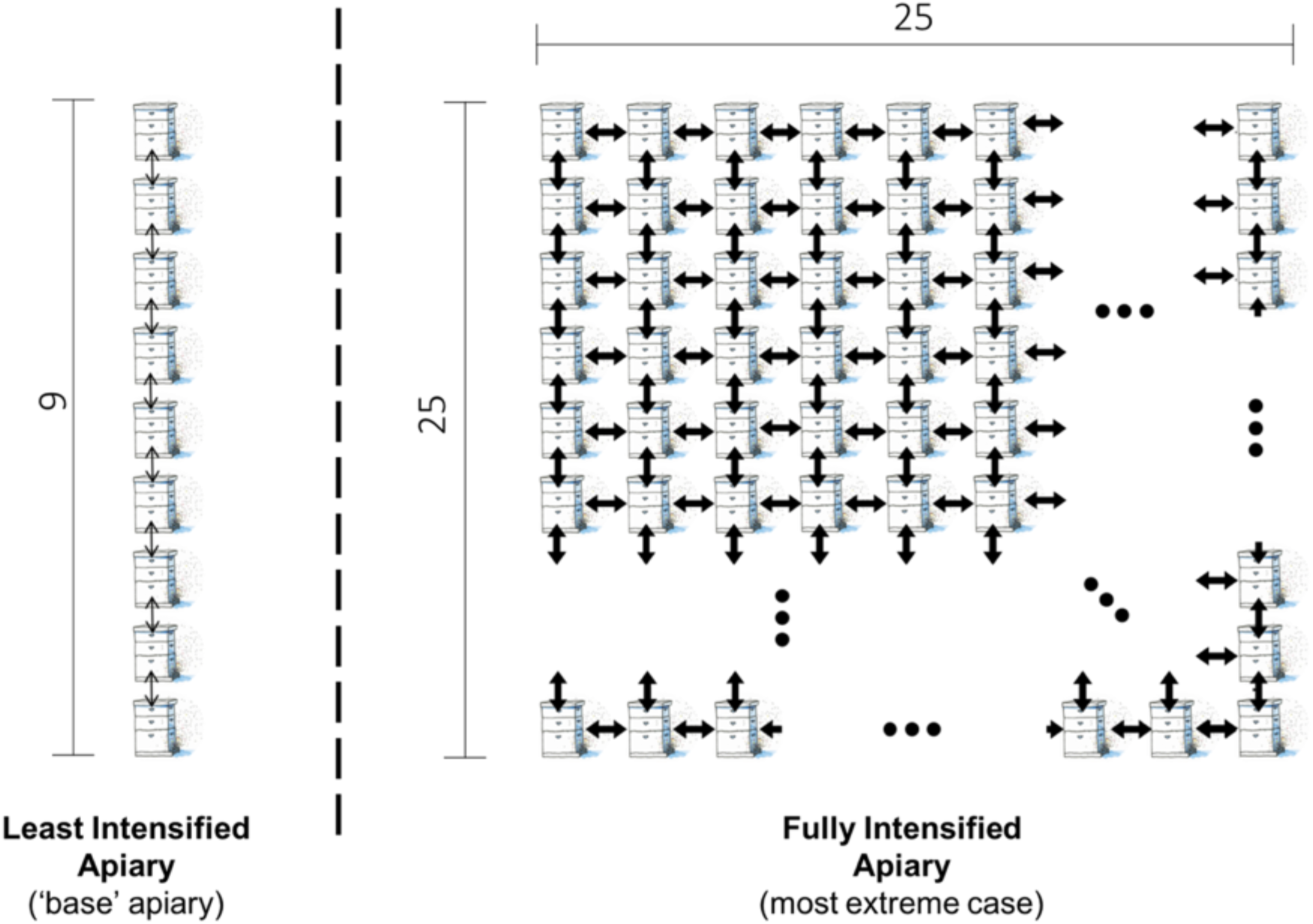
Illustrative schematic of ‘intensification’ as it is used in parts of this manuscript. We show the apiary used to calculate ‘base R_0_’ (left) compared to the intensified apiary (right) reflecting an increase in number of colonies from 9 to 225, a change from an array to a lattice, and a tenfold increase in movement of honeybees between colonies (illustrated using arrow weight) from a likelihood of 0.015 per bee per day to 0.15. Note that for the intensified apiary, not all 225 colonies are shown, with missing colonies denoted by ellipses (…).

We first derive a compartmental SI (Susceptible, Infected) model for pathogen transmission within an apiary. The model treats each colony as an individual population and allows for within colony as well as between-colony transmission (for nearest neighbours). Within a colony, honeybees are either susceptible to infection or infected (and infectious). We denote the number of susceptible honeybees in colony *i* at time *t* as *S_i_(t)*. Likewise, we denote the number of honeybees in colony *i* infected with the pathogen at time *t* as *I_i_(t)*. Susceptible honeybees in colony *i* become infected at rate β_ij_ following contact with an infected bee that resides in colony *j*. We assume that honeybees do not recover from infection. Honeybees are born at rate *ϕ*, have a natural mortality rate of *m*, and an additional mortality rate of *v* if infected. The following *2n* differential equations, [1], model disease transmission within and between *n* colonies in an apiary.

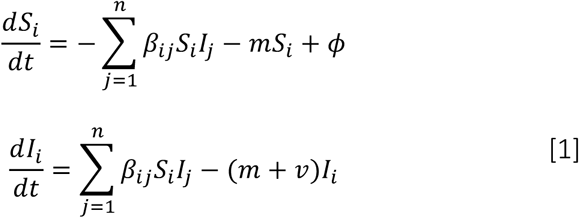

The matrix β=[β_ij_] will depend on the colony arrangement (see Fig. 1; and S.I. Section 1). The transmission rate between a susceptible and infected honeybee within the colony is *a*, and transmission between neighbouring colonies is *b*. We assume that honeybees are much more likely to become infected by a honeybee that resides within its home colony than by a honeybee from a neighbouring colony (i.e. *a*>>*b*). Note that for each apiary configuration to be possible and unique, the number of colonies (n) must be a perfect square, *n*=*L*^2^ where *L*≥3 (see Fig. 1). Therefore, the minimum number of colonies per apiary is 9, which has been observed to be the mean size of a hobbyist or small beekeeping operation (Mõtus et al., 2016; Pocol, Marghitas, & Popa, 2012). We complement our mathematical model [1] with the ABM; our ABMs are simulations of pathogen spread, through individual bee movements, across an apiary. Apiaries are differentiated by the same characteristics as in the mathematical model; a description of the ABM is available in the S.I. (Section 2). We use the ABM to make standalone predictions on the effects of different aspects of intensification on pathogen epidemiology (S.I. Figs. S3 & S4). We use the ABM to simulate disease dynamics for both different pathogen phenotypes (varying both pathogen virulence and transmissibility) and different apiary ecologies (varied as previously described in the number of colonies per apiary, layout, and likelihood of bees moving between colonies).

We can understand the dynamics presented by our models by focussing on the basic reproduction number, R_0_. R_0_ is a fundamental concept in infectious disease ecology, defined as the average number of secondary infections caused by one infectious individual in an otherwise entirely susceptible population (Anderson, May, & Anderson, 1992). We derive R_0_ expressions, using model [1], for each of the apiary configurations. R_0_ derivations using model [1] allow us to characterise the relationship between R_0_ and pathogen prevalence, defined as the proportion of honeybees within an apiary that are infected at the endemic equilibrium. For the ABM we calculate R_0_ values for particular parameter combinations by treating simulation outputs as ideal empirical data (Keeling & Rohani, 2008) and track the number of infections following the index case. The term ‘base R_0_’ is used throughout the remainder of this paper and refers to a value of R_0_ for a specific pathogen phenotype in a least intensified apiary (see Fig. 2). We determine how intensification affects R_0_ by separating R_0_ into a ‘base R_0_’ and an ‘additional R_0_’. The term ‘additional R_0_’ refers to the observed difference in R_0_ for a given pathogen phenotype when comparing a ‘lower intensity’ apiary to a ‘high intensity’ one (Fig. 2)

The most extreme plausible examples of intensification are used in these comparisons. Specifically, these are increases in colonies per apiary from 9 to 225 colonies, a change to a lattice configuration, and/or a tenfold increase in honeybee movement likelihood between colonies to 0.15 per bee per day, demonstrated in Fig. 2. Each is examined individually but we focus on the combined effect (reflected in Fig. 2). The difference in the R_0_ before and after intensification is how we calculate ‘additional R_0_’. This permits the interaction (non-additive) effects of our three aspects of intensification. The ‘additional R_0_’ can then be used in combination with the analytically derived relationship between R_0_ and prevalence (see model [1] & Results) to characterise how intensification affects disease prevalence. We focus on disease prevalence as both models show rapid pathogen spread across apiaries, such that infection prevalence at the endemic equilibrium was the major result differentiating modelling scenarios (S.I. Figs. S4 & S5).

## Results

The R_0_ expressions for apiaries with *n>1* colonies was calculated using the next generation method (van den Driessche & Watmough, 2002), (see S.I. Section 1).

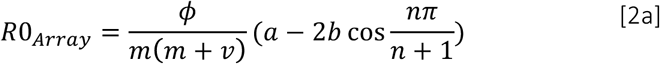

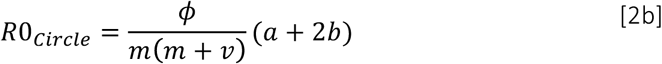

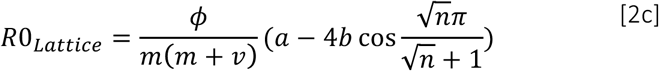

Both model [1] and the ABM simulations show that, for a given number of colonies per apiary, R_0_ is always greatest for the lattice arrangement — the most highly connected configuration. As the number of colonies per apiary increases (increasing n), the values of R_0_ in both the array and lattice configurations increase (Fig. 3a & 3b), while the R_0_ for the circular configuration remains unchanged (see R_0_ equations). The increase in R_0_ from the addition of colonies asymptotes quickly due to convergence in the mean number of neighbours across the apiary; this is also why the R_0_ for the circular apiary is independent of number of colonies as the number of neighbours per colony remains two.

**Figure 3:**
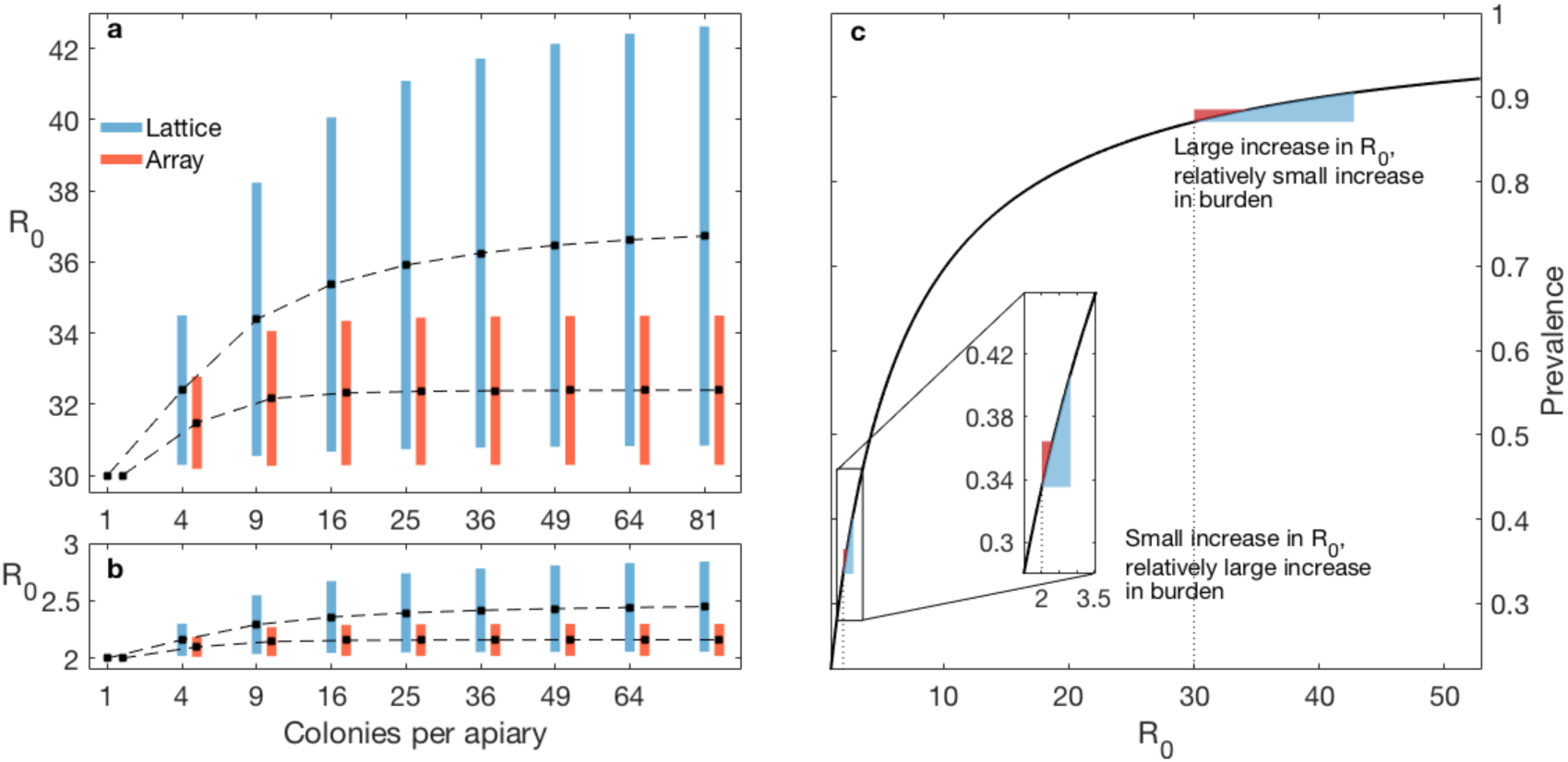
Relationships between number of colonies, R_0_, and prevalence. **a)** When R_0_=30 for a single colony-apiary, the addition of colonies yields a maximum increase in R_0_ of 12.7 for the lattice and 4.5 for the array. **b)** When R_0_=2 for a single colony, there is a maximum increase in R_0_ of 0.85 for the lattice and 0.29 for the array, when colonies are added. Recall that the R_0_ for the circle is independent of *n* (see [2b]), and hence absent from the figure. Black dots are values where between-colony transmission is held at 10% of total transmission, with the bottom and top of the bars representing 1% and 20% of the total transmission respectively. **c)** The relationship between R_0_ and disease prevalence. The range of R_0_ values is generated by varying the overall transmission rate (i.e. *a+b*) from 2.143×10^−6^ to 1.178×10^−4^ as reported by Roberts & Hughes (2015) for *Nosema ceranae*.

If R_0_>1, the pathogen will rapidly invade (see S.I. Section 1 &, Fig. S5) and each colony will reach a stable population size and infection prevalence, called the endemic equilibrium (See S.I. Section 1). Mathematically the disease prevalence at equilibrium for colony *j* is 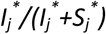, where 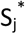 is the number of susceptible honeybees and 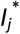 is the number of infectious honeybees in colony *j* at equilibrium. The endemic equilibrium for the circular configuration model can be solved explicitly (see S.I. Section 1). Due to symmetry, all colonies within the circular apiary have disease prevalence at the endemic equilibrium of:

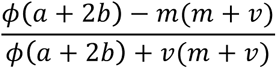

We can approximate the endemic equilibrium for the lattice and array configured models using perturbation theory, assuming 0 < *b* ≪ 1 (See S.I. Section 1). The approximate disease prevalence in colony *j* at equilibrium for a colony in the array or lattice configurations is:

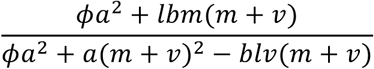

where *l* is the number of neighbours that colony *j* has. For any given set of parameters, we can therefore formulate both R_0_ and prevalence, allowing us to characterise the relationship shown in Fig. 3c.

We show analytically, and in the ABM (S.I. Section 3) that intensification in the form of an increase in colonies or an increase in movement between colonies increases R0 (Fig. 3a & 3b). Figure 4 shows the additional R_0_ caused by our most extreme plausible changes in apiary management. The change in R_0_ caused by increasing apiary size rapidly asymptotes (Fig. 3 a & b).

**Figure 4:**
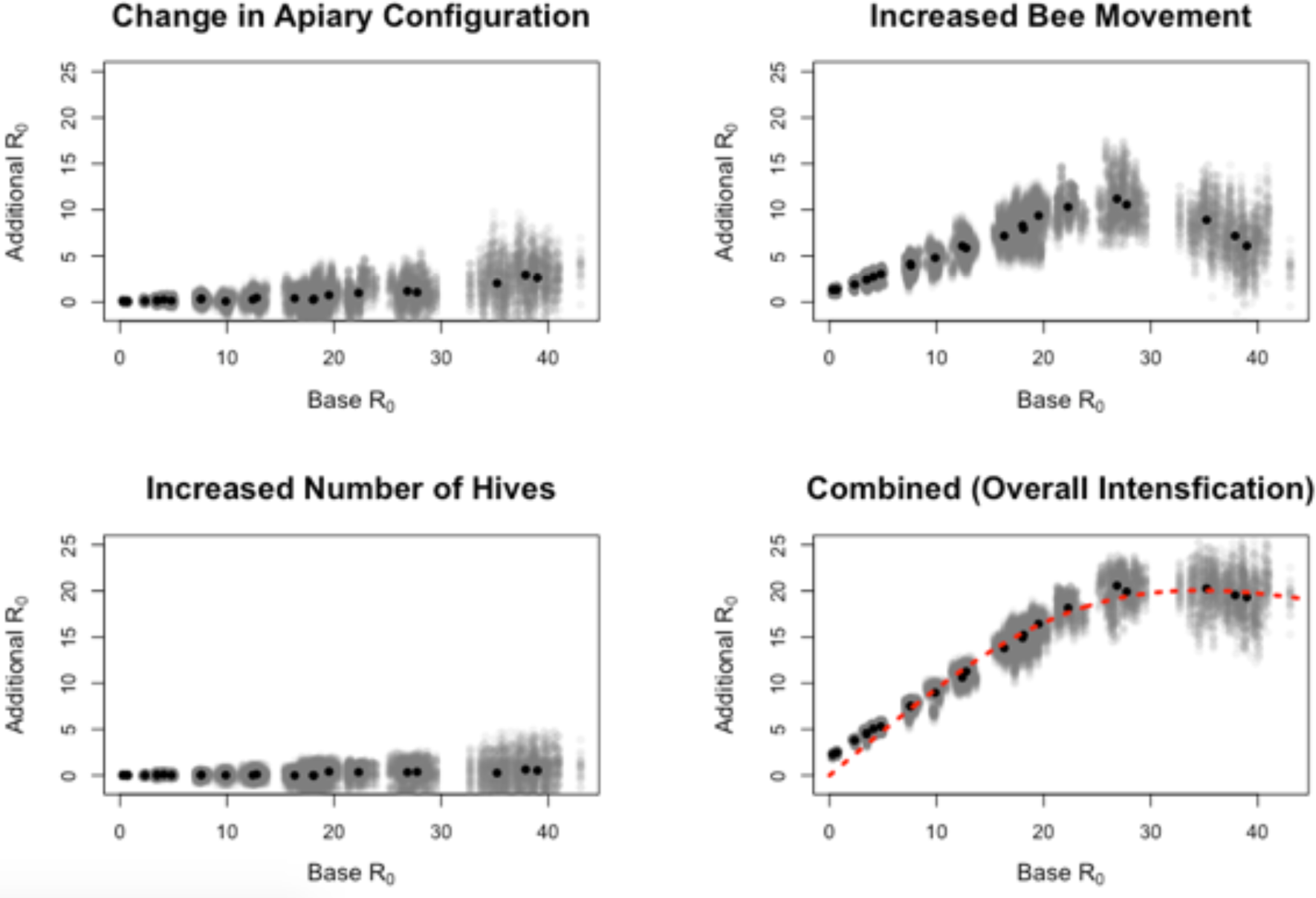
Simulation results from the ABM. The change in R_0_ caused by plausibly extreme increases in colonies per apiary, bee movement, and a change in configuration, across a range of different ‘base R_0_’ values determined by pathogen phenotype. Grey points represent individual simulation comparisons, black points represent mean values. Base R_0_ values are unevenly distributed across the range due to R_0_ being an emergent property of the system. We derive a non-linear relationship between ‘base R_0_’ and ‘additional R_0_’ for the ‘Combined’ treatment (represented in Fig. 2), plotted as a dashed red line.

Increasing movement between colonies has the strongest effect on R_0_ (Fig. 4). However, there are clear interaction effects present; the combined effect of all three aspects of intensification is greater than their additive sum. The effect of intensification is dependent on the base R_0_ – for small base R_0_, intensification causes little additional R_0_, but at intermediate or high base R_0_, intensification leads to large additional R_0_ (Fig. 4). The relationship shows a strong nonlinearity when examining all three aspects of intensification in combination.

By understanding the effect of intensification on R_0_ (Fig. 4) and by characterising the relationship between R_0_ and disease prevalence (Fig. 3c), we can show how intensification impacts disease prevalences. We approximate the non-linear relationship between ‘base R_0_’ (pathogen phenotype) and the ‘additional R_0_’ (effect of intensification) for the ‘Combined’ treatment (Fig. 4). We use a bootstrapping approach to create 1000 subsamples (subsample size = 10% of full sample with replacement) of our combined approach. Each subsample is used to generate a non-linear model of the form *y = ax/(b + x^c^)*, where *y* is ‘additional R_0_’ and *x* is ‘base R_0_’, using a nonlinear least squares approach in R (v 3.3.1). The relationship generated using the full sample is plotted in Fig. 4.

We combine this relationship characterising how base R_0_ affects intensified additional R_0_ (Fig. 4) with the derived relationship between R_0_ and pathogen prevalence shown in Fig. 3c, allowing us to predict how intensification impacts prevalences (Fig. 5). Fig. 5a shows the proportion of bees infected by a given (base R_0_) pathogen for the apiaries in Fig. 2. The difference in disease prevalence between these lines is the impact of intensification and is plotted in Fig. 5b. Fig. 5b shows a distinctly non-linear relationship between base R_0_ and the impact of intensification, with the impact of intensification peaking around base R_0_ = 3.3, and then rapidly declining. Even at its peak, the effect of intensification (which is as extreme as plausible), leads to an additional ~18% of bees infected at disease equilibrium.

**Figure 5:**
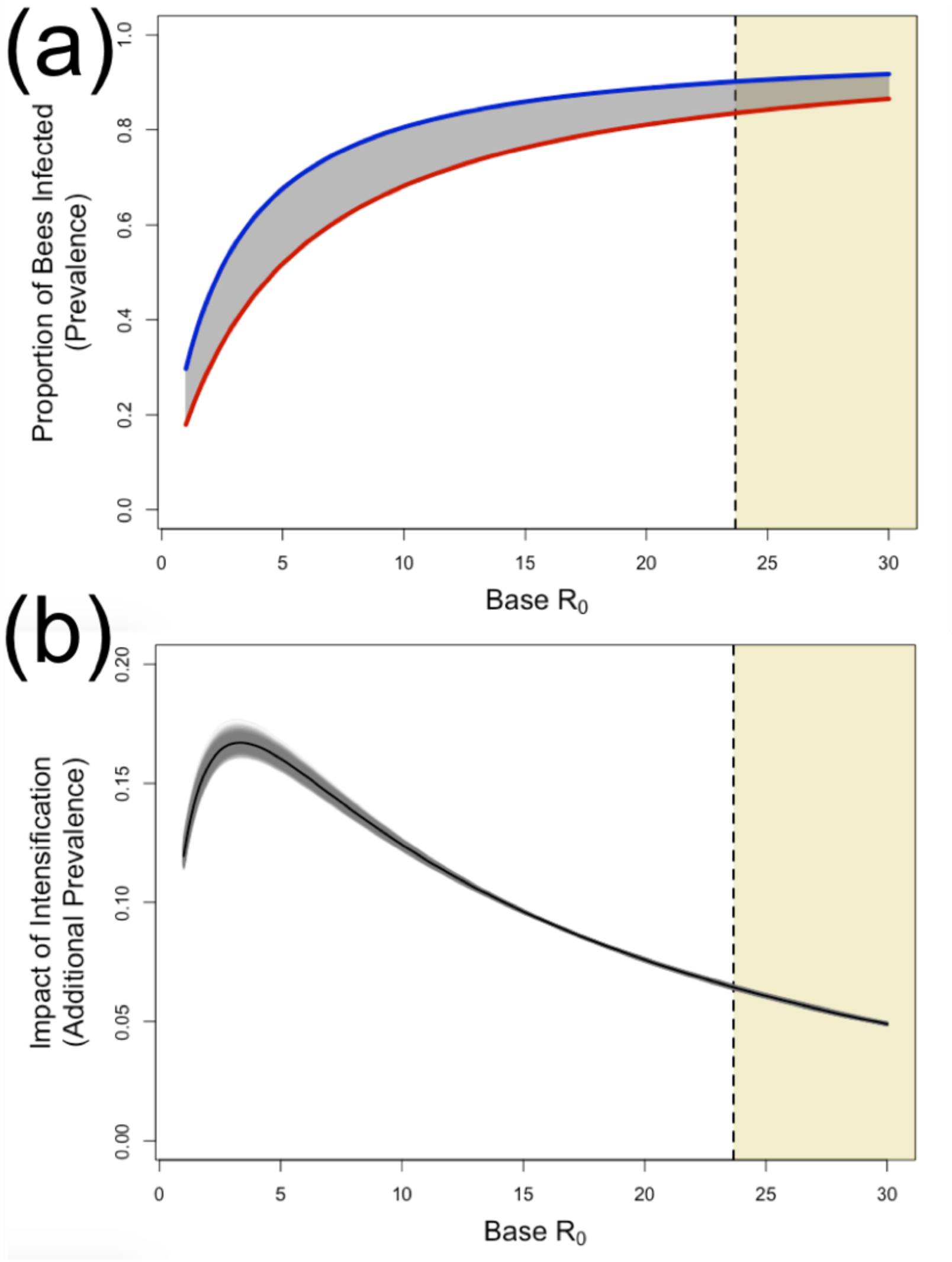
Consequences of intensification on disease prevalence for pathogens with different base R_0_ values, starting at R_0_ = 1.0008. Panel **(a)** shows the proportion of bees infected (prevalence) in non-intensified apiaries (lower red line) compared to intensified apiaries (upper blue line), calculated from the mean values derived in Fig. 4 and the relationship shown in Fig. 3c. The shaded grey area between these curves is the additional prevalence caused by intensification – the ‘impact of intensification’. This is plotted in panel **(b)** where the black line represents the mean relationship, and the grey lines represent 1000 bootstrapped samples. The vertical dashed line and yellow-shaded region of the graphs to the right of the dashed line show a lowest estimated value of R_0_ for *Nosema ceranae*.

We contextualize these results by calculating an estimate of the lower-bound of R_0_ value for a honeybee pathogen (see highlighted regions in Fig. 5). We identified this region based on empirical data for the microsporidian pathogen *Nosema ceranae;* this was the only pathogen for which experimentally derived transmission rates as well as robust information on mortality due to infection is available (Martín-Hernández et al., 2011; Paxton, Klee, Korpela, & Fries, 2007; Roberts & Hughes, 2015). To estimate the plausible R_0_ boundary in our model for this pathogen, we parameterised our mathematical model using the lowest empirically supported transmission value with the highest supported additional mortality, and fixed movement of honeybees between colonies at its lowest supported natural rate (Currie & Jay, 1991). We then calculated the R_0_ for a circular apiary due to its scale independence.

## Discussion

Our results present a counterintuitive picture of apicultural intensification and its consequences on disease prevalence within apiaries. Even in their most plausibly extreme cases, changes in the number of colonies, their spatial arrangement, and the movement of individual bees between colonies (reflecting intensification (Brosi et al., 2017)) had only a small effect on the severity of disease at the apiary level. Intensification leads to large gains in R_0_ when R_0_ is initially high and small gains in R_0_ when R_0_ is initially low (Fig. 4). However, increases in R_0_ cause large increases in prevalence only when R_0_ is initially low (Fig. 3c). Pathogens with a base R_0_ ≈ 3 benefit most from intensification in terms of increased prevalence (Fig. 5); however, the magnitude of this is moderate. As discussed below, we argue that there is likely to be a high base R_0_ in important honeybee diseases and therefore our models suggest that there is likely to be little effect of apiary-scale intensification on disease prevalence.

Our models most closely resemble the ecology of a directly transmitted microparasite able to infect individual honeybees at any life stage, conceptually similar to the microsporidian pathogen *Nosema* spp. (Fantham & Porter, 1912). *Nosema* is a major concern to beekeepers worldwide (Higes et al., 2008, 2009; Paxton, 2010), and has a minimum estimated base R_0_ of 23 (Fig. 5) when modelled here. We found that apicultural intensification, in the context of a pathogen with an initial R_0_ of 23, leads to a maximum 6.6% increase in disease prevalence. Our models predicted disease prevalences of up to 90% (Fig. 3, Fig. 5; S.I. Section 3), which while high, are empirically supported for the honeybee system (Higes et al., 2008; Kielmanowicz et al., 2015), and feature in other modelling studies that use similar transmission parameters to ours (Matt I. Betti, Wahl, & Zamir, 2014). *Nosema* was the only pathogen for which there are direct empirical studies characterising its transmissibility, however, other important honeybee pathogens are well studied, for example strains of deformed wing virus (DWV-B) (McMahon et al., 2016). While estimating an R_0_ for DWV-B is difficult due to active management by beekeepers, maximum reported prevalences that may be indicative of its true ‘unmanaged’ R_0_ are high, for example 73% in Natsopoulou et al. (2017) and 80% in Budge et al. (2015). These high prevalences are consistent with high R_0_ values (Fig. 3c & S.I. (Section 3)).

We additionally explored the behaviour of a more specific model, using an age-structured approach to infection dynamics, where only larvae are vulnerable to infection and develop into infectious adults with a high pathogen-associated mortality (as might be appropriate for pathogens such as the acute paralysis virus complex (Martin, 2001)), presented in the S.I. (Section 3). Convergence to equilibrium happens more slowly than the main model presented here, but still occurs quickly (within a single beekeeping season; see S.I. 3 Fig. S6). However adult-bee infection prevalence is far lower than seen in our SI model (S.I. Fig. S6) – this is in agreement with observations of lower prevalence of paralysis viruses (Budge et al., 2015). Notably, the endemic equilibrium prevalence increases only by small magnitudes as movement between colonies or apiary sizes are drastically increased (S.I. Fig. S6), in agreement with our main general result. This equivalence in behaviour between different models reflecting large disparities in infection mechanics, with empirically-supported different endemic prevalences, provides evidence that these results are likely generalisable to many honeybee pathogens.

We find rapid spread of a given pathogen across an apiary, which quickly reaches endemic equilibrium (S.I. Figs. S4 & S5). While pathogens with a higher R_0_ reach this equilibrium more quickly, there is universally rapid spread. Given this result, we focussed throughout this manuscript on the disease prevalence experienced at endemic equilibrium. This is important for our assumption that pathogen transmission (driven by movement of bees between colonies) only occurs between nearest neighbours. This assumption is conservative as rates of pathogen spread would be faster by virtue of not being limited to nearest-neighbour transmission. However, as we already observe rapid pathogen spread across apiaries, the effect of this conservative assumption should be negligible. The rate at which epidemics are established in our model is also in agreement with other honeybee pathogen models. For example, Jatulan, Rabajante, Banaay, Fajardo, & Jose (2015) show a single infectious adult causes an American Foulbrood (*Paenibacillus larvae*) epidemic that peaks within 50 days. Whilst they do not explicitly find an R_0_ for *P. larvae*, the short timescales characterising their epidemics are in line with ours (S.I. Section 3), suggesting high R_0_ values and that their model would behave similarly to ours at an apiary scale.

Changes in rates of bees moving between colonies emerged as a determining component of apicultural intensification (Fig. 4). One cause of this movement is honeybee drift (Jay, 1965) which can be managed through changes in the number of colonies and apiary configuration (Jay, 1966, 1968). Links between drift-mediated pathogen transmission and colony numbers have been documented for a variety of pathogens (Seeley & Smith, 2015) – including brood specialised and non-specialised, micro- and macro-parasites (Belloy et al., 2007; Budge et al., 2010; Dynes et al., 2017; Nolan & Delaplane, 2017). Larger numbers of colonies per apiary are a driver of higher drift (Currie & Jay, 1991), as are changes in apiary arrangement (Jay, 1966). This is why we focus on a ‘combined’ interpretation of intensification in this study (illustrated in Fig. 2), supported by our observation that changes in colonies per apiary and apiary size matter most when movement between colonies is high (Fig. 4; S.I. Fig. S4).

Our results contradict some empirical findings that larger apiaries are at higher risk of notably greater disease prevalences (Mõtus et al., 2016). However, our models do not account for landscape-scale movement of pathogens between apiaries. This is a phenomenon which has been well documented (Lindström, Korpela, & Fries, 2008; Nolan & Delaplane, 2017). Given our results, and empirical studies that did not find an association between colonies per apiary and disease risk (Giacobino et al., 2014), we argue that increasing the number of colonies in an apiary does not meaningfully alter within-apiary ecology to cause of increased disease prevalence. Larger apiaries may instead be more likely to import pathogens from other apiaries. Additionally, overstocking of colonies may lead to resource limitation and consequently impaired immune function (Al-Ghamdi, Adgaba, Getachew, & Tadesse, 2016; Pasquale et al., 2013). These effects are important for a broader understanding of honeybee epidemiology, but should be separated from the within-apiary processes studied here. Additionally, most honeybee infectious diseases are caused by multi-host pathogens shared with other wild bees (Fürst et al., 2014; Manley et al., 2015; McMahon et al., 2015, 2018). Honeybee colony density across a landscape therefore has implications for wild pollinator health (Cohen et al., 2017; Graystock et al., 2016), however our results suggest that increased stocking of honeybees may have smaller impacts on local pollinator infectious disease dynamics than may have been previously thought.

Two clear candidates for future development of this model include seasonality and demography, which are closely linked. Honeybee demography within a colony influences epidemiology (Betti, Wahl, & Zamir, 2016) due in part to the temporal polyethism of task allocation influencing exposure and immunity (Calderone & Page, 1996), as well as the flexible ability of honeybees to regain immune function when they revert roles (Amdam et al., 2005; Robinson, Page, Strambi, & Strambi, 1992)). However, patterns in how age and immunosenescence in honeybees relates to survival and infectiousness remain complicated (Roberts & Hughes, 2014). Analytically tractable models accounting for the role of this complex demography in understanding stress in a colony have only recently been developed (Booton, Iwasa, Marshall, & Childs, 2017), and extending these models to incorporate diseases at the apiary scale is challenging. However, notable phenomena worth pursuing include the role of male bees, which are known to be more easily infected, more infectious, and more likely to drift between colonies (Currie & Jay, 1991; Roberts & Hughes, 2015), as well as the role of robbing – where honeybees invade other colonies to steal food (Fries & Camazine, 2001).

Other industrialised agricultural livestock systems reflect extreme host densities similar to those in this study. However, the R_0_ for honeybee diseases may exceed that of other livestock diseases. We compare our lower threshold estimate for the R_0_ of *N. ceranae* to all available R_0_ values for livestock diseases that we could readily find in the literature (Fig. S8, see S.I. Section 4). Notably, all other livestock diseases for which R_0_ estimates exist show minimum R_0_ values far below our honeybee estimate, however examples of agricultural R_0_ values as high or higher than those we present for honeybees do also exist. There is therefore a clear need to develop explicit models of agricultural intensification scenarios for important agricultural disease.

Overall, our findings represent the first stage in developing robust epidemiological models for studying honeybee pathogens at an apiary scale. In the face of increasing challenges to global apiculture, our models predict that the size of apiaries *per se* is not causing notable increases in disease prevalence for important bee pathogens. Finally, this study demonstrates that conventional thought on how agricultural intensification influences disease may not be robust in the face of the

## Acknowledgements

LJB acknowledges funding from a Natural Environment Research Council training grant (NE/L002434/1). CR, MB, and AW acknowledge funding from a Biotechnology and Biological Sciences Research Council grant (BB/L010879/1). BJB, JCdR and KSD acknowledge funding from National Institutes of Health (R01-109501); the content of this study is solely the responsibility of the authors and does not necessarily represent the official views of the National Institutes of Health.

## Authorship

All authors contributed to conceptualisation and scope definition of the study. LJB, CR, MB developed approach. Mathematical modelling was undertaken by CR, AW, and MB. Computational modelling by LJB, KD, and MB. Model scope and parameterisation by LJB, KD, JCdR, BJB, LW. LJB and CR created figures, interpreted results and drafted manuscript with guidance and input from all authors. All authors contributed to further drafting, revision, and finalisation.

